# A natural history model for planning prostate cancer testing: calibration and validation using Swedish registry data

**DOI:** 10.1101/402743

**Authors:** Andreas Karlsson, Alexandra Jauhiainen, Roman Gulati, Martin Eklund, Henrik Grönberg, Ruth Etzioni, Mark Clements

## Abstract

Recent prostate cancer screening trials have given conflicting results and it is unclear how to reduce prostate cancer mortality while minimising overdiagnosis and overtreatment. Prostate cancer testing is a partially observable process, and planning for testing requires either extrapolation from randomised controlled trials or, more flexibly, modelling of the cancer natural history.

An existing US prostate cancer natural history model (Gulati et al, Biostatistics 2010;11:707-719) did not model for differences in survival between Gleason 6 and 7 cancers and predicted too few Gleason 7 cancers for contemporary Sweden. We re-implemented and re-calibrated the US model to Sweden. We extended the model to more finely describe the disease states, their time to biopsy-detectable cancer and prostate cancer survival. We first calibrated the model to the incidence rate ratio observed in the European Randomised Study of Screening for Prostate Cancer (ERSPC) together with age-specific cancer staging observed in the Stockholm PSA (prostate-specific antigen) and Biopsy Register; we then calibrated age-specific survival by disease states under contemporary testing and treatment using the Swedish National Prostate Cancer Register.

After calibration, we were able to closely match observed prostate cancer incidence trends in Sweden. Assuming that patients detected at an earlier stage by screening receive a commensurate survival improvement, we find that the calibrated model replicates the observed mortality reduction in a simulation of ERSPC.

Using the resulting model, we predicted incidence and mortality following the introduction of regular testing. Compared with a model of the current testing pattern, organised 8 yearly testing for men aged 55–69 years was predicted to reduce prostate cancer incidence by 0.11% with no increase in the mortality rate. The model is open source and suitable for planning for effective prostate cancer screening into the future.

**Author summary:** A naïve perspective is that cancer screening is simple: people are screened, some cancers are detected early, and cancer mortality rates decline. However, the mathematics for screening becomes difficult quickly, it is hard to infer causation from observational data, and even large randomised screening studies provide limited evidence. Simulations are therefore important for planning cancer screening.

We found an older US prostate cancer natural history model to be poorly suited for contemporary Sweden. We therefore re-implemented and re-calibrated the US model using data from Swedish registries.

Our revised model, the Stockholm “Prostata” model, provides predictions similar to those observed in the detailed Swedish registers on prostate cancer incidence and mortality. By modelling the mechanisms of the screening effect, we can predict the benefits and harms under a range of screening interventions.

## Introduction

Cancer screening policies must balance the benefits and potential harms based on uncertain and incomplete evidence. It is difficult to infer causation from observational data, and even large randomised screening studies provide limited evidence. Simulations using natural history models can provide further insights. Such natural history models describe the course of the disease from onset to progression through to death. Calibration of such models against observed disease incidence patterns with and without screening can be used to improve our understanding of the mechanisms for disease progression and cancer screening interventions. Simulations of the natural history of disease can be used to bring together evidence from specific randomised controlled trials with data from other sources and to generalise the results from specific population structures and disease prevalence [1, 2]. Finally, these simulations can also be used as a basis for cost-effectiveness analysis in order to make informed decisions on cancer screening interventions [3–5].

Our application relates to prostate cancer. The prostate is a male reproductive organ that, together with other glands, is responsible for the production of semen. As men age, they are more likely to suffer from prostate enlargement or prostate cancer. In Sweden, prostate cancer accounts for a third of male cancer diagnoses and a fifth of male cancer deaths [6].

Evidence from ERSPC suggests that PSA testing can reduce prostate cancer mortality by approximately 20% over 13 years [7]. There are two other large randomised studies of prostate cancer screening: the Prostate, Lung, Colorectal and Ovarian Cancer Screening Trial did not find any significant reduction in prostate cancer mortality when the control arm included high levels of opportunistic PSA testing [8]. The recent Cluster Randomized Trial of PSA Testing for Prostate Cancer found that with 40% attending the clinical visit in the screening arm for a single PSA screen in men aged 50–69 years lead to an estimated mortality reduction of 4% at ten years [9]. Although PSA testing is common in many western countries, testing is not systematically organised, and the balance of harms versus benefits of PSA testing is uncertain [4].

PSA testing in Sweden continues to be common and new prostate cancer tests are becoming available. To assess whether organised prostate cancer testing would be beneficial, we sought to develop a well-validated prostate cancer simulation model for Sweden.

There are few existing models for the natural history of prostate cancer. However in order to make full use of the detailed longitudinal Swedish registers and allow the natural history model to represent the mortality rate ratio observed in the ERSPC [7], we adapted and extended an existing model [10–12] with a more detailed natural history. This is important for modelling risk-stratified prostate cancer testing in combination with new screening tests.

Our objectives are to describe a contemporary, validated prostate cancer screening model and to apply that model to predict key screening outcomes under different PSA screening scenarios for Sweden. The longer-term goal is to use this simulation model to plan for better prostate cancer testing and screening.

## Results

### Model overview

We adapted a prostate cancer screening model from the Fred Hutchinson Cancer Research Center (FHCRC) [10–12]. The FHCRC model simulated for individual life histories, coupling PSA trajectories with the disease onset and progression from localised to advanced disease by Gleason score (low-moderate versus high grade). The FHCRC model used inputs from the Prostate Cancer Prevention Trial (PCPT) [13] and US PSA test patterns [14], and was calibrated to US data before and after the introduction of PSA testing, and validated against (a) the prostate cancer mortality RR from the ERSPC screening trial and (b) US prostate cancer incidence. Starting with the model from [12], we re-implemented and extended the model to include additional states by T-stage/M-stage and Gleason scoring as shown in Fig 1 (more recent developments of the FHCRC model are described in the Discussion). We also used more detailed inputs for calibration and validation. Data sources for the model inputs, calibration and validation included: the National Patient Register (including data on cancer treatment), National Prostate Register (cancer incidence by Gleason score, T-stage and M-stage), Total Population Register (defining the at-risk population), Cause of Death Register, the Stockholm PSA and Biopsy Register (SPBR), and the PCBaSe research database for prostate cancer survival. Details are provided in the Materials and Methods.

**Fig 1.**
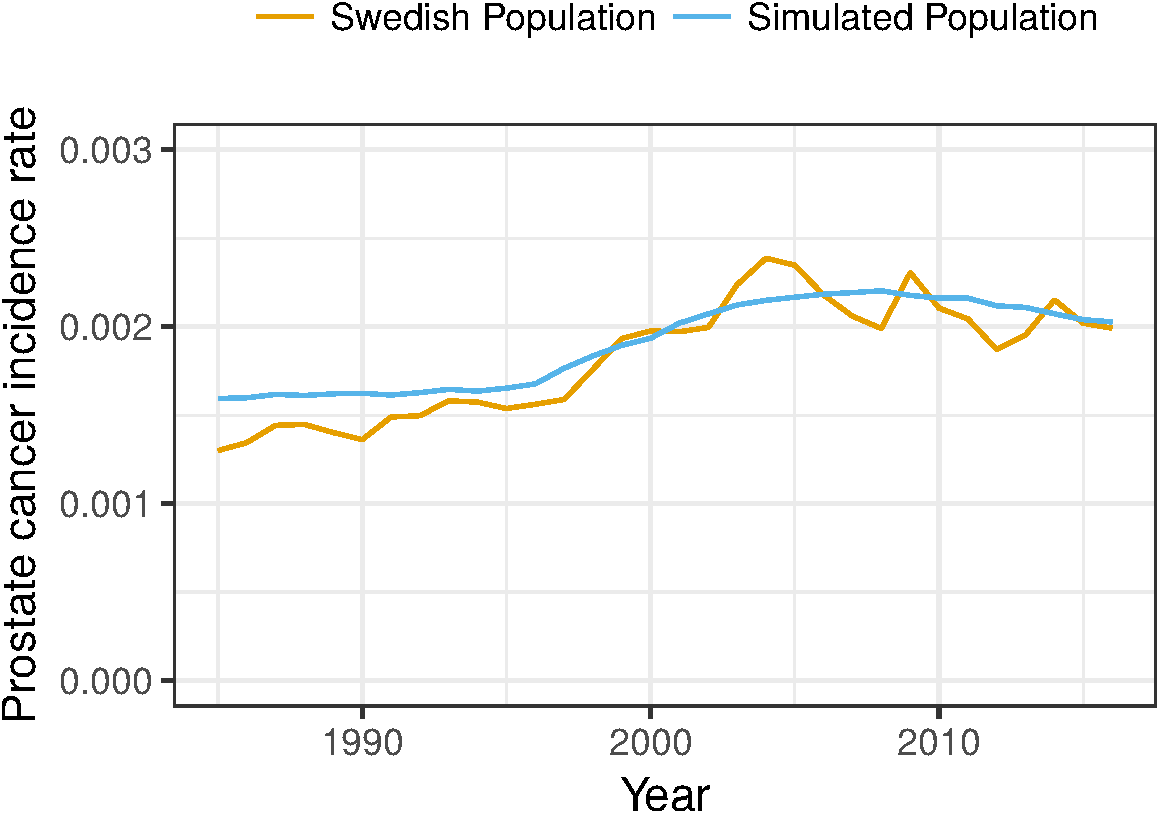
Schematic of the prostate cancer natural history model reflecting disease onset, progression and survival in the absence of screening. Individuals are assumed to be healthy at age 35 years; they may progress to preclinical cancer states with fixed Gleason score, but with progression by T-stage and to metastatic cancer; preclinical cancers may be clinically diagnosed from nine different states, with survival from prostate cancer death modelled from the time of clinical diagnosis; death due to other causes is represented as a competing event.

### Model calibration

Model calibration was undertaken in two steps. First, we calibrated for the relative distributions of incident cancers from contemporary Sweden and the screening effect on incidence from the ERSPC. Second, we calibrated to survival from contemporary Sweden. The Swedish calibration targets were from a PSA tested population. We modelled for PSA testing in this population using data from the SPBR (details are provided in PSA testing sub-model). The initial uptake was modelled by age and calendar period and PSA re-testing was modelled by age and PSA-value; PSA testing rates by age and calendar period are shown in Fig 2. The probability of having a biopsy following a positive PSA test (i.e. biopsy compliance) was modelled by age and PSA value using the SPBR (see Table 2 in the S1 Appendix).

**Fig 2.**
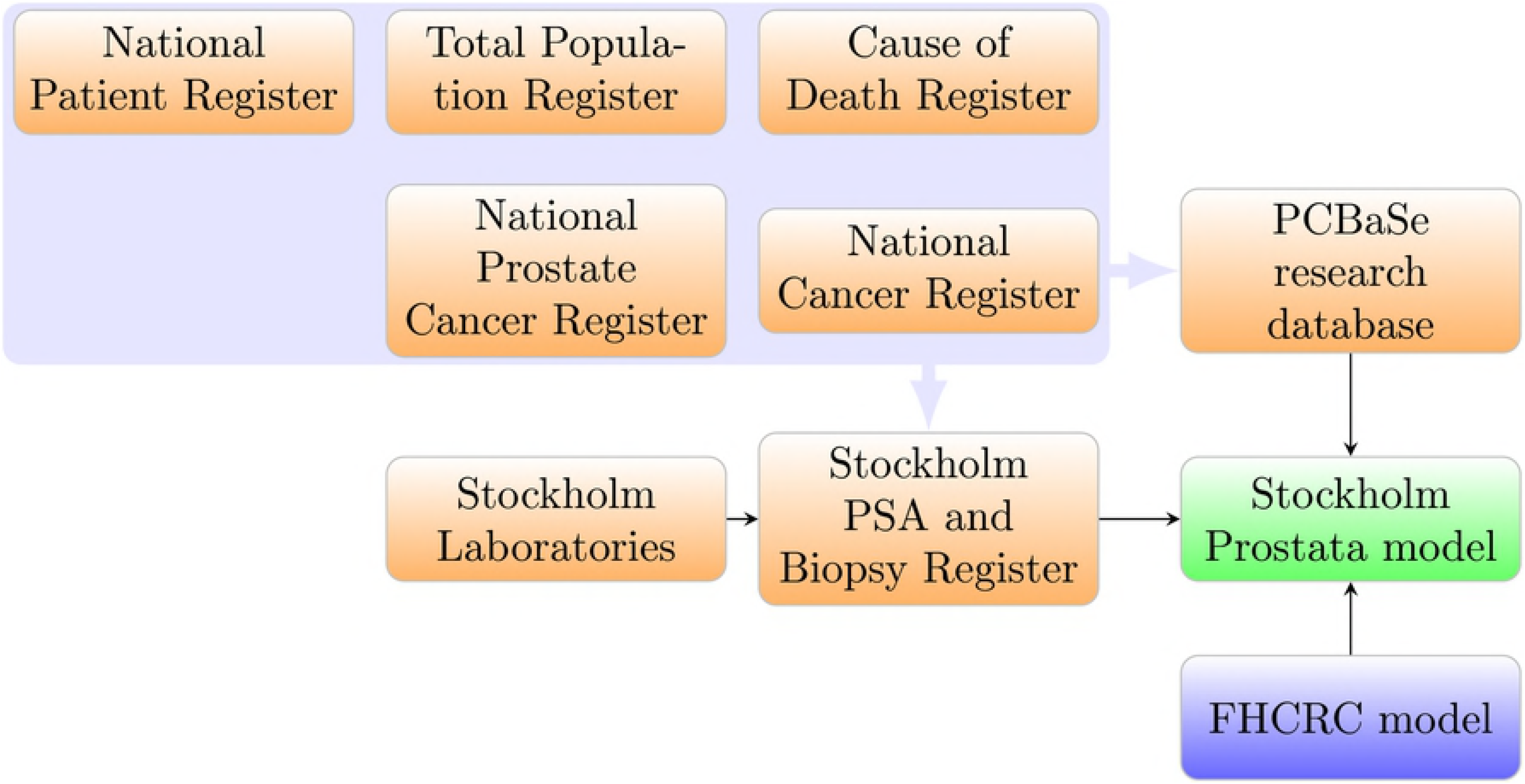
Modelled current PSA testing rates per person-year for ages 40-80 years and the calendar period 1995-2014 for men without an existing prostate cancer diagnosis. The white contour lines indicates the rates 0.1, 0.2 and 0.3. The modelled values are based on data from the Stockholm PSA and Biopsy Register [15].

#### Calibrating to the relative distributions of cancer staging

The observed relative distributions of incident cancer stages at diagnoses were used as calibration targets for modelling several prostate cancer natural history parameters (Tables 1 and 2). Importantly, we modelled for transitions between T-stages and fitted the relative distributions for the Gleason scores and cancer T and M stages by age groups; see Fig 3. We included different T-stages to support more detailed modelling of treatment and survival. The use of relative proportions allows for the absolute incidence rates to be used for validation. The calibration used a reconstruction of a contemporary Swedish population with data on PSA test uptake, health state proportions at diagnosis, and survival from a screened population. A total of 4392 diagnoses in the ages 50–74 from a three-years interval (2011–2013) were used as calibration targets.

**Table 1.** Population characteristics used to adapt the model to the Swedish context.

| Parameter | Description | Source |
| --- | --- | --- |
| External inputs | PSA trajectories | FHCRC [11] |
|  | Natural history prior to diagnosis | FHCRC [11, 12] |
| Directly observed | Treatment | SPBR |
|  | PSA and biopsy compliance | SPBR |
|  | PSA re-testing | SPBR |
| Calibrated parameters | PSA test rates | SPBR |
|  | Survival | PCBaSe [17] |
|  | Gleason distribution | SPBR |
|  | Progression from T1–T2 to T3–T4 | SPBR |
|  | Biopsy undetectable in T1–T2 | ERSPC (incidence RR) [7] |
| Validation targets | Cancer incidence (screened & unscreened) | Stockholm & Sweden [30] |
|  | Cancer mortality (screened & unscreened) | Stockholm & Sweden [30] |
|  | Screening mortality rate ratio | ERSPC (mortality RR) [7] |

**Table 2.**
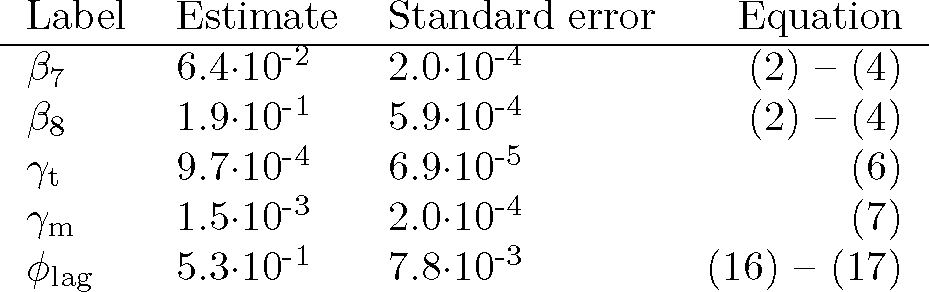
Estimated parameters from calibration procedure.

| Label | Estimate | Standard error | Equation |
| --- | --- | --- | --- |
| $\beta_7$ | $6.4 \cdot 10^{-2}$ | $2.0 \cdot 10^{-4}$ | (2) – (4) |
| $\beta_8$ | $1.9 \cdot 10^{-1}$ | $5.9 \cdot 10^{-4}$ | (2) – (4) |
| $\gamma_t$ | $9.7 \cdot 10^{-4}$ | $6.9 \cdot 10^{-5}$ | (6) |
| $\gamma_m$ | $1.5 \cdot 10^{-3}$ | $2.0 \cdot 10^{-4}$ | (7) |
| $\phi_{lag}$ | $5.3 \cdot 10^{-1}$ | $7.8 \cdot 10^{-3}$ | (16) – (17) |

**Fig 3.**
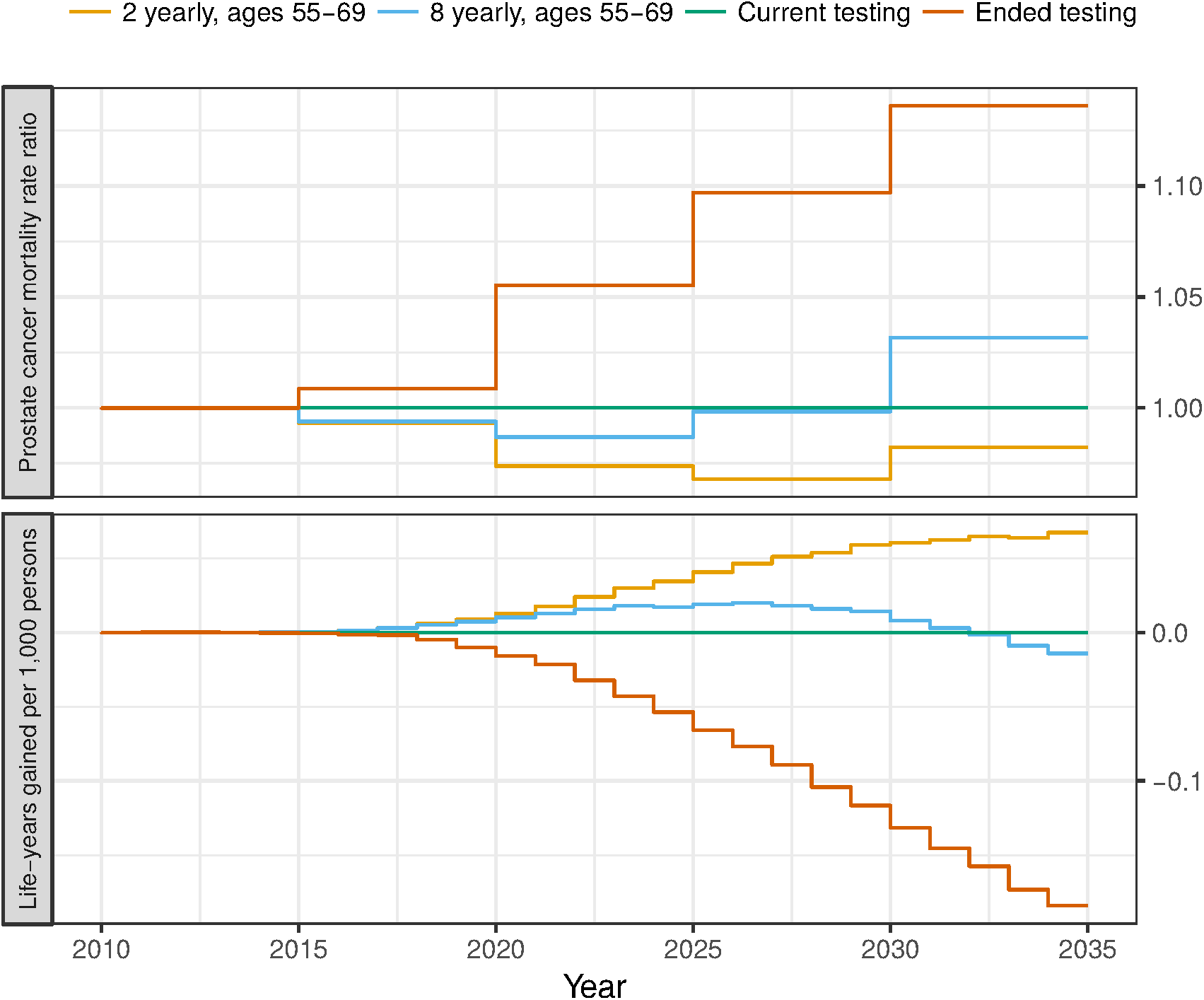
Fitted Gleason, T-stage and metastatic proportions of cases in 2011-2013 by age groups to that observed in the SPBR register.

#### Calibration of screening effects on incidence

In addition to the calibration of the screened Swedish population, we also simulated for the screened and unscreened arms from the ERSPC to replicate results from the 13 years of follow-up [7]; for details on the reconstruction of the ERSPC trial, see Materials and Methods. The ERSPC rate ratio of prostate cancer incidence (1.57, 95% confidence interval (CI) 1.51–1.62) was used in the calibration, while the ERSPC mortality RR of prostate cancer was used for validation. To calibrate to the ERSPC incidence rate ratio and to model indirectly for tumour size, we introduced a parameter for the proportion of the time from onset that a T1–T2 cancer would not be biopsy detectable [16]; we estimated that the T1–T2 cancers would on average be undetectable at biopsy for 47% of the time before they progressed to T3–T4 cancers.

#### Calibrating to survival from diagnosis

To calibrate prostate cancer survival, we compared simulated survival from a contemporary Swedish population, including longitudinal screening and treatment patterns, with observed 10- and 15-year survival from the PCBaSe database [17, 18]. The PCBaSe database contained 93014 men from the Swedish National Prostate Cancer Register diagnosed in the period 1998–2014 linked with the health and population registers. Prostate cancer survival was stratified by M stage, Gleason score (≤ 6, 7, ≥ 8), PSA (< 10, ≥ 10) and ten-year age groups. Predictions from the calibrated model are displayed as Kaplan-Meier survival curves and the observed 10- and 15-year survival are displayed as point estimates with 95% CIs in Fig 4. The calibrated model has a clear separation in survival for men diagnosed with either Gleason 6 or Gleason 7 cancers; see Fig 2 in the S1 Appendix for a comparison with the FHCRC model, where survival is similar for Gleason 6 and 7 cancers. We did not model for survival by treatment, as we did not have detailed information on the reasons for treatment assignment.

**Fig 4.**
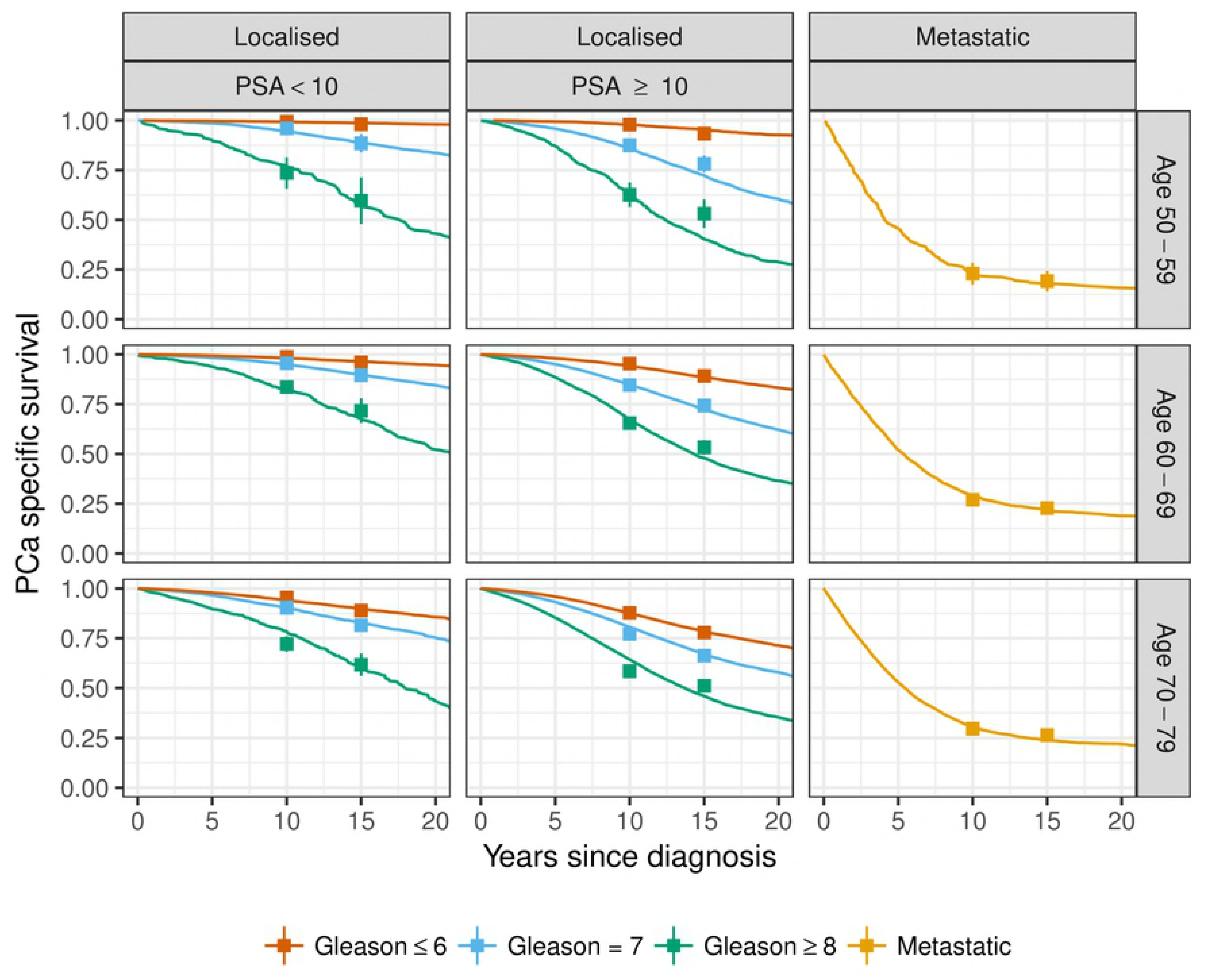
Simulated survival from the calibrated model displayed as Kaplan-Meier survival curves together with the observed 10- and 15-year survival from PCBaSe displayed as point estimates with 95% confidence intervals. Survival is stratified by age at diagnosis, PSA at diagnosis, Gleason score and cancer extent.

### Model validation

#### Population prostate cancer incidence

In Fig 5, we compared the age-standardised prostate cancer incidence rates from the simulation with that of Sweden during 1985–2016 which included the introduction of PSA testing. There is evidence for a good fit although the rapid increase in incidence following the introduction of PSA testing was not fully captured. This over-smoothing is possibly due to the PSA uptake sub-model having few degrees of freedom. Importantly, this is a validation and we did *not* calibrate for prostate cancer incidence rates.

**Fig 5.**
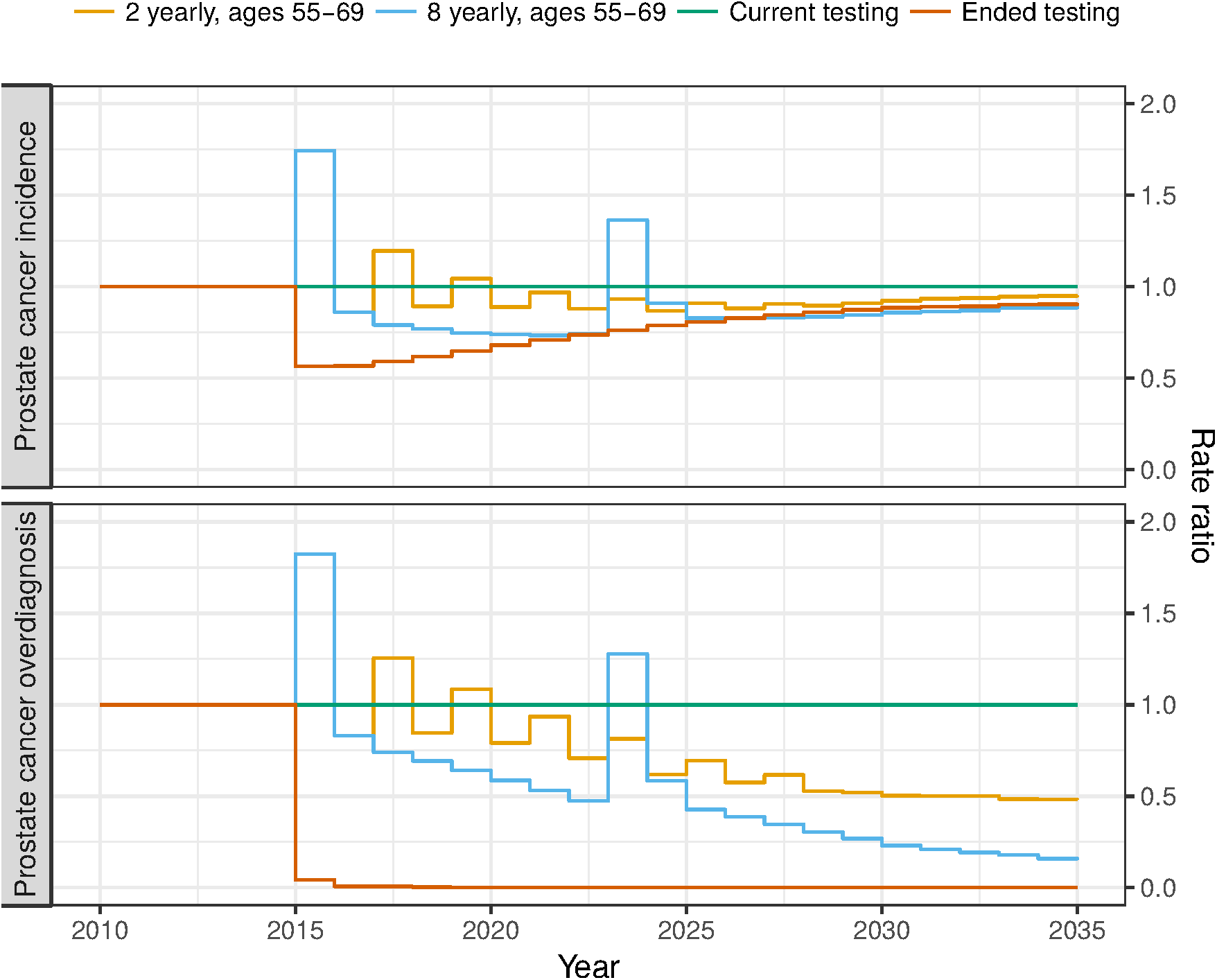
Age-standardised prostate cancer incidence rates from the simulation model compared with those observed in Sweden.

#### Screening effect on mortality

In contrast to the incidence increase from ERSPC, which was used for calibration, the mortality decrease was used for validation. Using simulations of the screened and unscreened arms in ERSPC (see Materials and Methods), we estimated the 13-year mortality rate ratio using Poisson regression. The ERSPC reported a mortality RR of 0.79 (95% CI 0.61–0.88), whereas our calibrated model predicted a RR of 0.784 (95% Monte Carlo interval (MCI) 0.781–0.786).

### Model predictions

When planning for prostate cancer testing policies, the following measures were considered to represent the burden of disease: prostate cancer incidence rate; prostate cancer overdiagnosis rate, where overdiagnosis is defined as the lifetime risk of having a prostate cancer diagnosis that would never have been clinically detected prior to death due to another cause; prostate cancer mortality rate; and life expectancy. We predicted these measures for a policy that replaces the current testing pattern (see Fig 2) with regular prostate cancer testing during ages 55–69 years: regular testing was introduced from 2015 at age 55 years for those born in 1960 and in later birth cohorts. Using this policy introduction, regular testing had completely replaced current testing across ages 55–69 years after 15 years. Our modelling of organised screening only specifically addresses the effect of screening intensity for the targeted age groups.

In Fig 6, we predicted prostate cancer incidence and overdiagnosis rate ratios for 20 years of 2-yearly testing, 8-yearly testing and the complete cessation of asymptomatic testing in comparison with the current testing pattern. The 2-yearly testing scenario resulted in a minor reduction, RR 0.98 (95% MCI 0.98–0.98), in prostate cancer incidence and a larger decrease in prostate cancer overdiagnosis, RR 0.80 (95% MCI 0.79–0.80), over 20 years compared with the current testing pattern. The less intensive 8-yearly testing scenario substantially reduced the prostate cancer incidence, RR 0.90 (95% MCI 0.89–0.90), and the reduction of prostate cancer overdiagnosis was even larger, RR 0.57 (95% MCI 0.56–0.57), compared with the current testing pattern. The hypothetical cessation of all PSA testing for asymptomatic men in 2015 would result in a substantial decrease, RR 0.74 (95% MCI 0.74–0.75), of prostate cancer incidence compared with the current testing rates over 20 years as well as no overdiagnosis of asymptomatic men.

**Fig 6.**
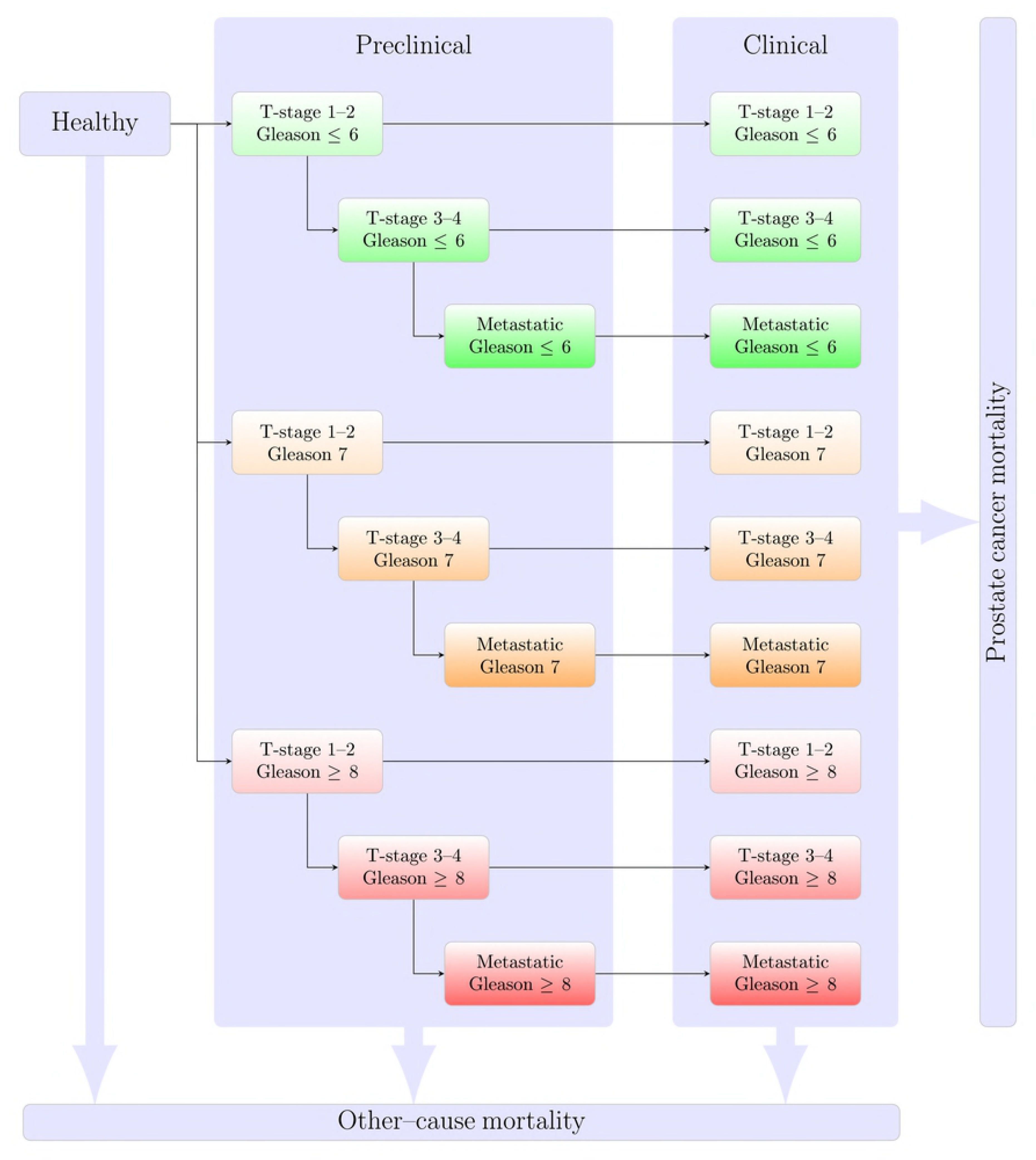
Predicting incidence and overdiagnosis rate ratios for 2-yearly and 8-yearly screening between 55 and 69 years of age and the cessation of asymptomatic testing compared with current testing uptake. The changes in testing policy were introduced in 2015 for a population reflecting the Swedish age-structure.

The purpose of early detection for prostate cancer is to lower prostate cancer mortality and increase the life expectancy. To assess these effects, we predicted mortality rates and life-years gained for the different PSA testing policies (Fig 7). Both the mortality rates and the life-years gained were expressed relative to the current PSA testing pattern. We predict that the broad introduction across the 1946–1965 birth cohorts contributed to the mortality reduction, which, while wearing off towards the end of the 20 year period, causes an increase in mortality, particularly for the 8 yearly testing. The relative effect on the mortality is considerably smaller than the effect on the incidence and while the 2 yearly testing pattern slightly reduces mortality, RR 0.98 (95% MCI 0.97–0.98), the 8 yearly testing pattern has a similar mortality, RR 1.01 (95% MCI 1.00–1.02), as the current uptake pattern for the predicted 20 years. Similarly the 2 yearly testing pattern slightly increased the life expectancy, 0.03 (95% MCI 0.03–0.04) life-years gained per 1,000 persons, and the 8 yearly testing pattern did not noticeably affect the life expectancy, 0.01 (95% MCI 0.00–0.01) life-years gained per 1,000 persons, compared to the current uptake pattern for the predicted 20 years. The hypothetical scenario of cessation of PSA testing for asymptomatic men in 2015 was predicted to significantly increase prostate cancer mortality over 20 years, RR 1.08 (95% MCI 1.07–1.08), and reduce the life expectancy by −0.07 (95% MCI −0.09—0.05) per 1000 persons. The effect is smaller than the 20% prostate cancer mortality reduction observed in the ERSPC study as the current PSA uptake pattern is less intensive than ERSPC and the lower biopsy compliance observed in Sweden (see Table 2 in the S1 Appendix).

**Fig 7.**
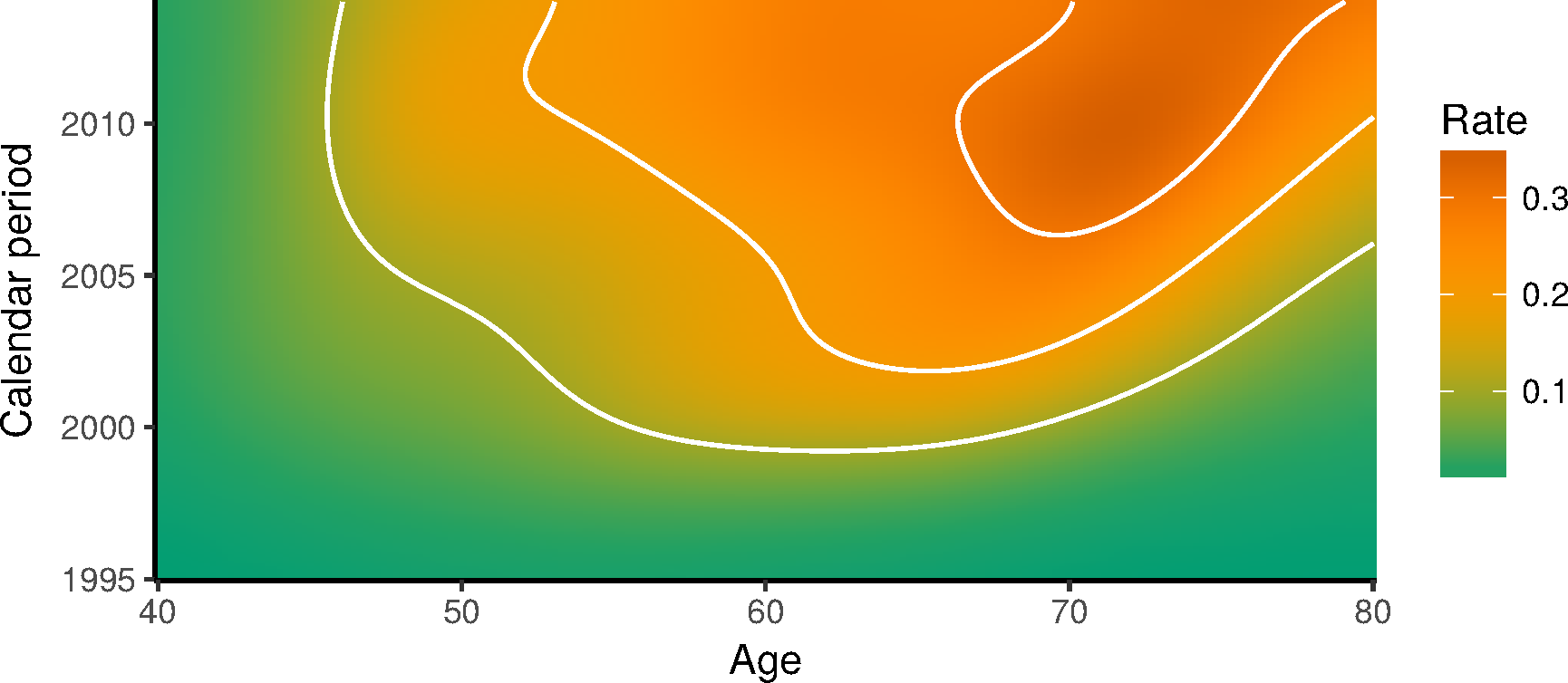
Predicting mortality RRs and life-years gained for 2-yearly and 8-yearly screening between 55 and 69 years of age and the cessation of asymptomatic PSA testing compared with current testing patterns. The shifts in testing policy was introduced 2015 on a population reflecting the Swedish age-structure.

The modest mortality reductions are potentially explained by relatively high levels of testing under the current PSA testing, and the use of the currently observed biopsy compliance for all predicted scenarios. These reductions are also comparable to the non-significant mortality reduction found in the Prostate, Lung, Colorectal and Ovarian (PLCO) cancer screening trial [8], where there were high levels of PSA testing in the control arm [19].

## Discussion

Our aim was to develop, calibrate and validate a prostate cancer natural history model that could be used to evaluate prostate cancer testing. Using extensive Swedish data resources, we extended an older US-calibrated prostate cancer natural history model for the Swedish population and validated the new model. We then used the revised model to predict longer-term patterns of prostate cancer incidence and mortality in Sweden.

One of the challenges with natural history models is finding a model that is biologically meaningful and representative, whilst being mathematically simple and potentially estimable. Another challenge is that many of the parameters of a cancer’s natural history are either not observable, such as the initial onset of disease, or are only partially observed at specific time points, such as the size of a tumour at the time of diagnosis.

Investigators are divided in how to resolve these challenges. One school uses very simple models with expert judgement for the effectiveness of interventions. The validity of the predictions depend on the accuracy of the experts. A second school uses Markov models fitted to evidence from randomised controlled trials (RCTs) to assess the effectiveness of specific interventions within the follow-up from the RCTs. The validity is limited by the available RCT evidence, with strong limitations for predicting outside of the observed data. A third school uses more detailed natural history models and simulate for individuals. The validity of the predictions primarily reflects the validity of the natural history model. We are firmly in the last of these three schools. We have previously modelled cancer screening using both simple and more complex Markov models, and found issues with validity for the simple models and issues with model complexity for scaling more detailed Markov models to combinations of natural history and test states by time in state [20].

One potential criticism of many microsimulation models for cancer screening is that their complexity is coupled with a lack of model detail and that the source is usually closed. The US-funded CISNET collaboration has provided detailed model documentation [21] and some models (e.g. FHCRC) are available on request. We addressed this criticism by making all of our code open source and easily available (https://github.com/mclements/microsimulation and https://github.com/mclements/prostata). We encourage other microsimulation modellers to make their code openly available, which will lower the entry requirements for other investigators. If the cost of entry remains high, then a closed source consulting model will continue to be predominant.

There are several potential limitations. First, the revised natural history model was less accurate for modelling event rates at older ages (e.g. over 80 years of age; see S1 Appendix). This is consistent with observations that Nordic prostate cancer mortality rates are typically higher than rates in US populations. We suggest caution when interpreting incidence for Nordic populations for several reasons: the higher Nordic rates may lead to greater absolute declines in rates, leading to more effective screening; and the point estimate for the mortality reduction due to screening was higher in the Göteborg site, although there was no statistically significant heterogeneity between the ERSPC sites [7, *p* = 0.4]. More accurate modelling at older ages would require a more detailed natural history model. Second, it is difficult to assess whether the natural history model is causal and accurate: the disease process is only partially observed and the biology represented using a simple mathematical representation. Third, the prediction of age-standardised mortality rates were slightly lower than that observed in the Swedish population. This underestimation could be due to e.g. changes in Gleason grading, where the Gleason and T-stage distribution at diagnosis was based on data from patients diagnosed 2011–2013 whereas the survival by stage was based on patients diagnosed 1998–2014.

Finally, as individuals were followed for up to 15 years, the survival calibration was influenced by earlier Gleason grading practices, possibly leading to an overestimation of the risks for Gleason*≤*6 cancers. Nonetheless, we expect that our predictions will have strong internal validity, as the simulations allow for carefully controlled experimental conditions.

Strengths of our approach include the wealth of detailed longitudinal data available from Sweden, and that we have made the model open source. Our natural history model can support an evidence-based approach to assessing whether the introduction of organised re-testing or screening would be effective and cost-effective. The Swedish Prostata model was branched from the FHCRC prostate cancer model in 2013. Since that time, both the Swedish Prostata model and the FHCRC model have incorporated a number of similar extensions, including T-stage development and more detailed modelling of Gleason grading [5]. A key difference is that the updated FHCRC model includes a two-parameter model for cancer onset [22]. We are currently investigating whether to incorporate these extensions into the Swedish Prostata model. A key advantage of the Swedish Prostata model is the availability of detailed longitudinal data on PSA values and prostate biopsies linked with clinical outcomes. The US CISNET prostate cancer models have historically relied heavily on un-linked, cross-sectional SEER data. In contrast the Swedish high coverage registry data is well suited for modelling the disease progression and treatment pathways within men, and would potentially improve the model validity.

Our choice of modelling approach included model calibration for some key parameters in both unscreened and screened populations (Table 1). To assess whether the adapted model was valid for Sweden, we compared the model predictions with observed population incidence. This approach demonstrates both the strengths and potential weaknesses of our model.

Our model is now well suited to the health economic evaluation of new prostate cancer screening tests. In particular, we have modelled for Gleason ≤ 6 cancers, which typically have very good prognosis, from Gleason 7 or Gleason ≥ 8 cancers, where the last category has particularly poor prognosis. The new prostate cancer tests have focused on maintaining sensitivity for more aggressive prostate cancers, such as Gleason 7 or higher, with reductions in the incidence of small Gleason ≤ 6 cancers and negative biopsies.

From the section on Model predictions, we found evidence to suggest that organised screening would reduce overdiagnosis without increasing mortality compared to current screening practices. Future work is needed to investigate refined screening strategies and evaluation of cost-effectiveness.

## Acknowledgement

We are grateful to Pär Stattin and Fredrik Sandin for providing data from PCBaSe, and for data management of the Stockholm PSA and Prostate Biopsy Register by Pouran Almstedt and the late Peter Olausson. We thank the following for discussions: Erwin Laure, Jan Adolfsson, Markus Aly, Tobias Nordström and þorgerður Pálsdóttir.

## Materials and Methods

In this section, we will describe the various data sources used to develop the model, explain the model formulation, outline the methods for the calibration and validation of the model, and finish with a description of the model implementation.

### Data sources

We have integrated multiple sources of data in order to extract relevant Swedish prostate cancer statistics for our model. The linkage between the different sources is illustrated in Fig 8. Detailed individual data on men who had a PSA test or a prostate core biopsy were extracted from the Stockholm PSA and Biopsy Register. Using the unique Swedish personal identification number [23], we linked the study cohort to a number of population registers, including the National Cancer Register (NCR) and the National Prostate Cancer Register (NPCR). The NCR included data on tumour extent (loco-regional vs distant) and the date of diagnosis; the NPCR contained additional clinical information, including Gleason score and TNM cancer stage classification. The NCR and NPCR were known to cover 96.3% and 94% of cancer patients, respectively [18, 24]. A dynamic cohort (with men moving in and out of the Stockholm County) was defined via the Population Register which contains information on migrations within Sweden as well as external migration.

**Fig 8.**
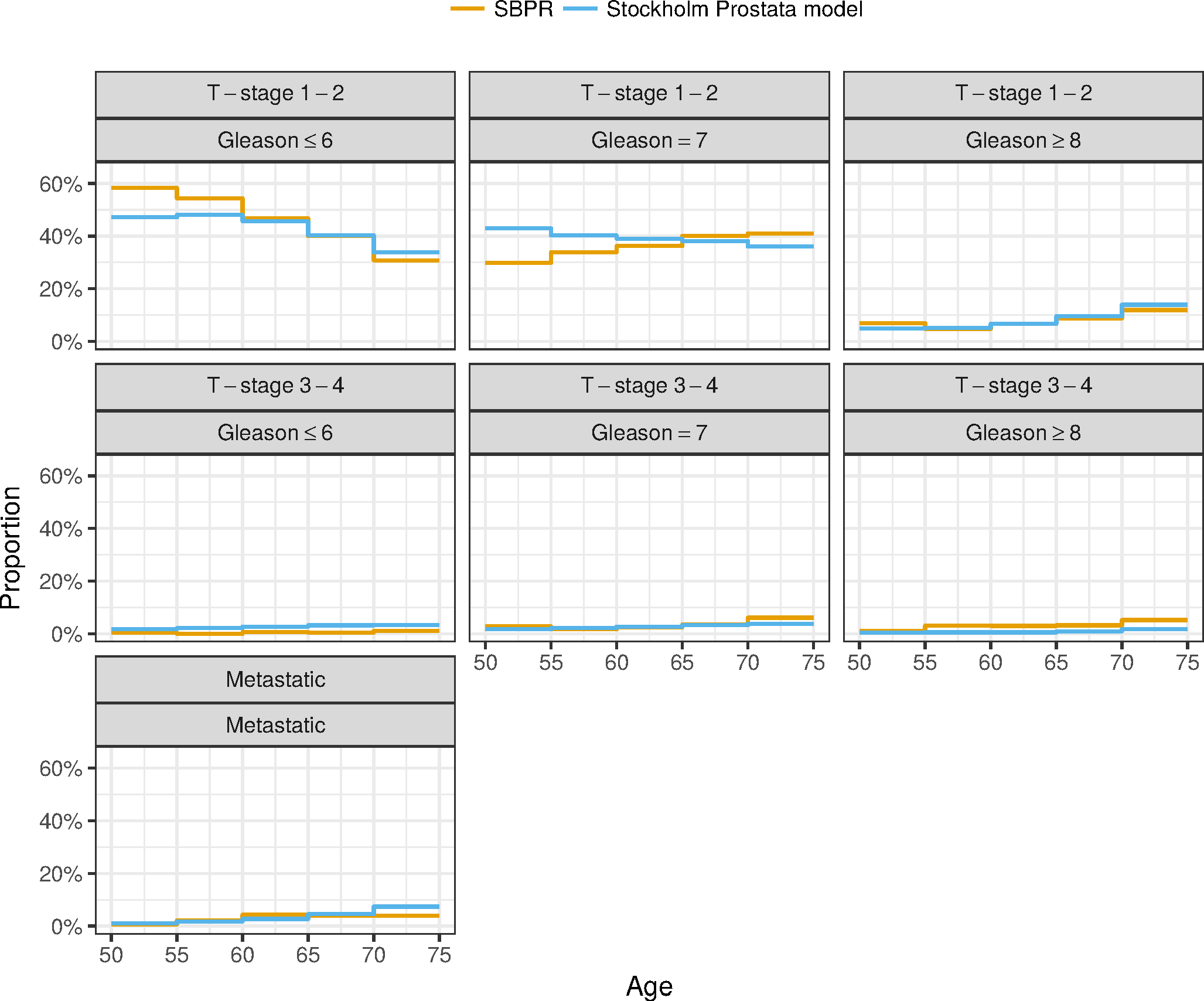
Overview of data sources and their linkage.

From PCBaSe, a research database linking the NPCR with other health and population registers, we extracted survival following a prostate cancer diagnosis by calendar period, age, stage, PSA value and Gleason score; and treatment modality by calendar period, age and Gleason score.

Data on prostate cancer incidence by calendar year, age, stage and Gleason scores were extracted from the NCR and the NPCR. Prostate cancer mortality by calendar year and age were obtained from the Cause of Death Register. The Total Population Register was used to calculate the male population at risk by calendar period and age.

### Model description

The prostate cancer natural history model links PSA growth with prostate cancer progression.

The two main components of the model are: (i) longitudinal PSA growth; and (ii) transitions between natural history disease states, as shown in Fig 1. The PSA growth is expressed functionally in Equation (1) and the transitions between states is defined in equations (5) – (11). Cancer onset is assumed to be independent of PSA, and that PSA rises faster after cancer onset. The distribution of Gleason score is assumed to be multinomial with the proportions modelled as a function of age at cancer onset, with no de-differentiation (or change in Gleason score) after onset.

#### PSA growth

The change in PSA values after cancer onset is assumed to differ by Gleason score, with a specific change for Gleason score 6 and below (G6-), Gleason 7 (G7) and Gleason 8 and above (G8+).

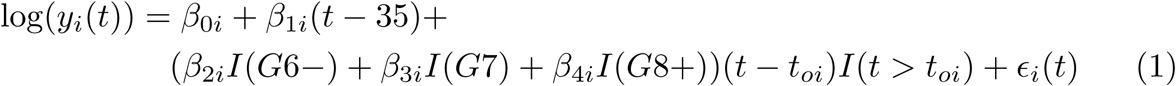

where *y*_*i*_(*t*) is measured PSA at age *t* for subject *i, I*(*A*) is a 1 if *A* is true and 0 otherwise, *t*_*oi*_ is the age of cancer onset and where

- 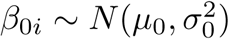, is a random intercept
- 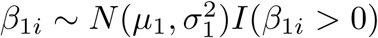, which is a random slope
- 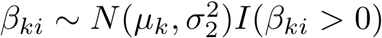, *k* = 2, 3, 4, is a Gleason-specific random change of slope after cancer onset
- *ϵ*_*i*_(*t*) *∼ N* (0, *ϕ*^2^), represents measurement error.

The random slopes N(*µ,σ*^2^)I(*β*>0) are truncated distributions to ensure that PSA growth is monotonically increasing.

The values for *µ*_1_, *σ*_1_, *µ*_4_ and *σ*_4_ were from [11], while the estimates for *µ*_2_, *σ*_2_, *µ*_3_ and *σ*_3_ were weighted sums to separate the estimates of Gleason *≤* 7 from [11] into Gleason *≤* 6 and Gleason 7.

#### Gleason score distribution

The Gleason score assigned to an individual at cancer onset is dependent on the age at cancer onset according to the probabilities modelled via the multinomial logistic regression in Equations (2) – (4) as illustrated in Fig 9 (right panel).

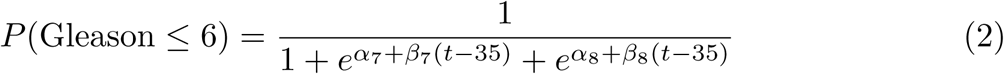

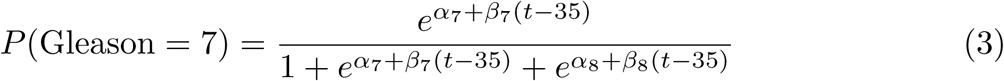

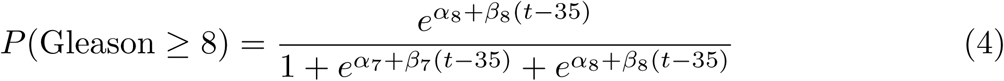

where *t ≥* 35.

**Fig 9.**
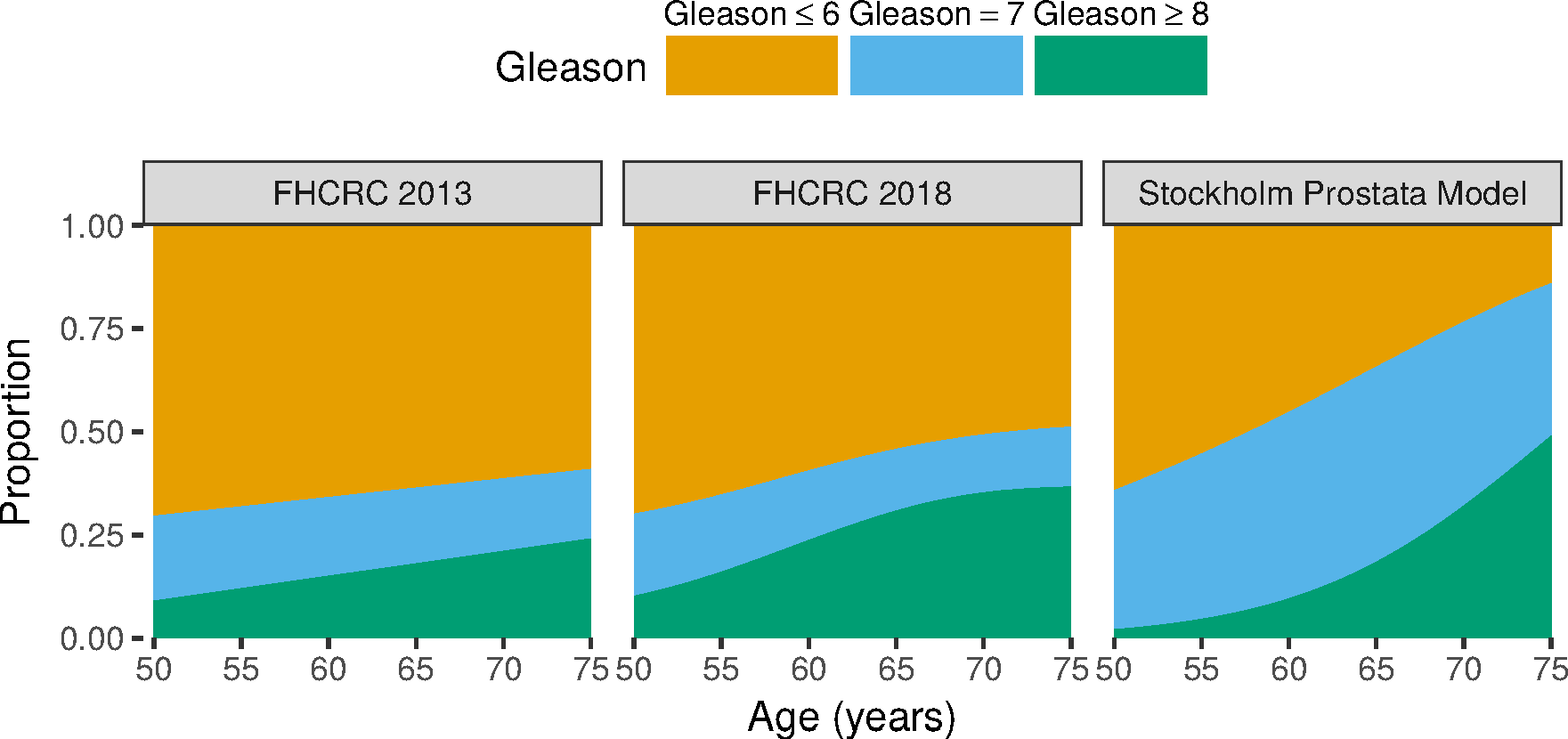
Comparing the modelled proportion of Gleason scores at cancer onset from FHCRC model in 2013 and 2018 with the Stockholm Prostata model.

#### Transition rates

Transitions between different states in the model (i.e. healthy, localized states, metastatic states and death) are simulated via events, which occur with different rates.

The disease onset (a transition from the healthy to a localized state) is modelled via a time-dependent hazard (from age 35) as

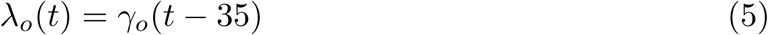

which means that the time-to-event follows a Weibull distribution (shape parameter 2 and scale parameter 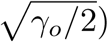. The cumulative distribution (the complement of the survival function) for the time to cancer onset is hence 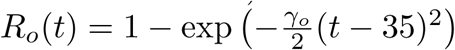 [11]. The event density, which is simply the probability density for the Weibull distribution with parameters as above, represents the rate of cancer onset per unit time (see Fig 1 in the S1 Appendix).

Transitions between disease states are dependent on age (*t*) and the individual log-PSA-values 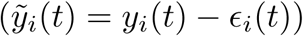. The model includes T-stage transitions within localised cancer states for preclinical cancer. The transition from T1–T2 to T3–T4 is the same for all Gleason categories and is described in Equation (6). *γ*_*t*_ is the hazard of transitioning to T3–T4 and the time-dependence comes from the log-PSA levels.

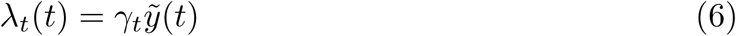

The rate from T3–T4 to metastatic disease is proportional to PSA and *γ*_*m*_ which is the metastasis hazard (see Equation (7)). Note that the FHCRC model used *γ*_*t*_ to represent the parameter for the transition rate from onset to metastatic [11].

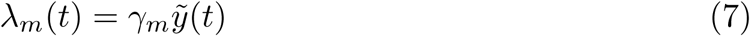

The clinical diagnosis rate for localised cancer onset for Gleason score 7 and lower (Equation (8)) and Gleason score 8 and higher (Equation (9)) are proportional to PSA and 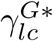, which is the clinical diagnosis hazard for localised cancer for the two Gleason score categories. As per the older US model, we combined the Gleason ≤6 and 7 scores for these transitions due to a lack of informative data.

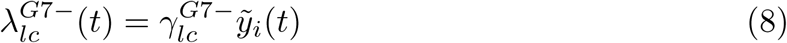

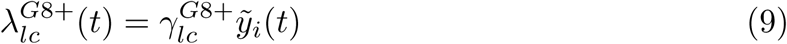

The rate to clinical diagnosis after metastatic onset for Gleason score 7 and below (Equation (10)) and Gleason score eight and above (Equation (11)), is proportional to PSA and 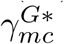 is the post-metastasis clinical diagnosis hazard for the two Gleason score categories.

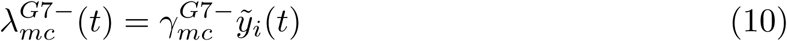

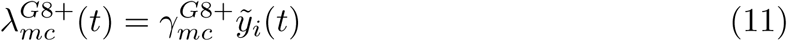

#### PSA testing sub-model

Diffusion of a new health technology into a population is a dynamic process. This process may reach a stationary state after a longer period of time For PSA testing, test uptake was distributed across a range of ages over a comparatively short period, such that the PSA test patterns varied substantially by birth cohorts. PSA test uptake is required for calibrating the model to screened populations.

The natural history model is calibrated to data that are observed both before and since the introduction of PSA testing. In particular, we have survival data for men diagnosed for prostate cancer from 1998, which is after the introduction of PSA testing This requires that we accurately model for PSA uptake and re-testing and for treatment to represent the men at risk for prostate cancer incidence, survival and mortality.

The PSA sub-model represents uptake of the PSA test together with the pattern of PSA re-testing. Uptake was modelled as: (i) a function of age for cohorts born from 1960; (ii) a function of calendar period multiplied by a factor for birth cohort for birth cohorts born before 1932; and (iii) a mixture of (i) and (ii) for the birth cohorts between 1932 and 1960. Mathematically, age-specific uptake (i) is modelled by the cumulative density function for a log-logistic cure model, such that

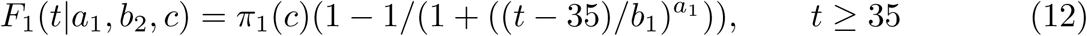

where *t* is age at uptake, *π*_1_ is the proportion of men ever having a PSA test, *c* is the calendar year of birth (or birth cohort), and where *a*_1_ and *b*_1_ are the shape and scale for a log-logistic distribution for those men who ever have a PSA test. The calendar-specific uptake for the older cohorts (ii) is modelled by

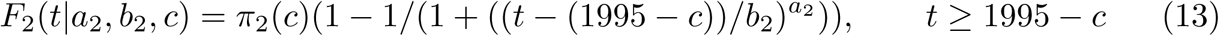

where *π*_2_ is the proportion of men who ever have a PSA test, and where *a*_1_ and *b*_1_ are the shape and scale for a log-logistic distribution for those men who ever have a PSA test. Finally, for the intermediate birth cohorts, *t*_1_ is sampled from *F*_1_, *t*_2_ is sampled from *F*_2_, and *t*_1_ is selected over *t*_2_ with probability (1960 −*c*)*/*(1960 – 1932).

PSA re-testing is modelled using a Weibull cure model, such that

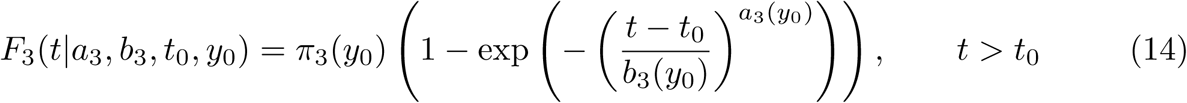

where *t*_0_ is the age at the previous PSA test, *y*_0_ is the value of the previous PSA test, *π*_3_ is the proportion of men who will ever have a re-test, and where *a*_3_ and *b*_3_ are the shape and scale parameters of the Weibull distribution for those who ever have a re-test.

For re-testing, the parameters *π, a*_3_ and *b*_3_ were estimated using a Weibull cure model stratified by the five-year age groups (30–34, 35–39,…, 85–89, 90+) and by PSA values ([0, 1), [1, 3), [3, 10), [10, *∞*)) at the previous PSA test. The parameters *π*_1_, *π*_2_, *a*_1_, *a*_2_, *b*_1_ and *b*_2_ were calibrated to observed PSA test rates for Stockholm using a Poisson likelihood.

#### Biopsy sensitivity and compliance

For men who had a PSA value above 3 ng/mL, the proportion complying with a subsequent biopsy varied by PSA values and age, and was estimated from the SPBR (see Table 2 in the S1 Appendix). We also modelled for whether a prostate cancer was biopsy-detectable, assuming that a cancer was not initially detectable for a proportion *ϕ*_lag_ (16) – (17) of the time from cancer onset to the development of a T3–T4 cancer. Our approach varied from Wever et al. 2010 [16], who modelled for the sensitivity of a PSA test to detect a cancer by stage, irrespective of the time from cancer onset. The probability of a biopsy (Bx) rendering a diagnosis (Dx) depends on the biopsy sensitivity, the biopsy compliance and the probability of cancer:

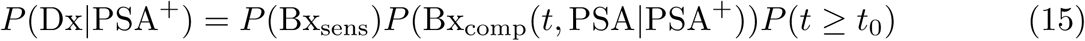

where P(t ≥ t_0_) is the probability of having had a cancer onset, *P* (Bx_comp_(*t*, PSA | PSA^+^)) is probability of performing a biopsy after a positive PSA test depending on age and PSA value and *P* (*Bx*_sens_) is the biopsy sensitivity as expressed below:

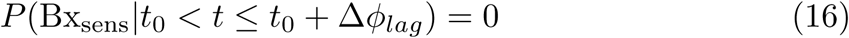

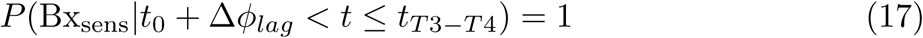

where Δ = *t*_*T* 3_–_*T* 4_ – *t*_0_ is the time with a T1–T2 cancer.

#### Treatment sub-model

Probabilities for treatment assignment to either active surveillance, radical prostatectomy, radiotherapy or androgen deprivation therapy were assessed from the SBPR. These values were stratified by five year age groups and Gleason score (see Fig 3 in the S1 Appendix).

#### Survival sub-model

Survival from cancer diagnosis to death due to screening was calibrated to the NPCR. In summary, we first simulated for Sweden for 1998–2014 including screening uptake with survival distributions from SEER (localised vs metatstatic), which are also used as inputs to the FHCRC model. NPCR survival estimates were available at ten and fifteen years after diagnosis by age groups for the period 1998–2014; for non-metastatic cancer, survival was available by Gleason score and for PSA less than 10 ng/mL and for 10 ng/mL and over. We compared the Kaplan-Meier estimates of survival from diagnosis from the simulated data with observed Kaplan-Meier estimates for men diagnosed with prostate cancer in Sweden. We did not calibrate to observed survival from the pre-PSA era, as such estimates were not available.

One significant modelling challenge is selecting and fitting a mathematical representation for the effect of cancer screening. For cancers with a short period between a screen-detected diagnosis and a counter-factual clinical diagnosis, a common model is to represent differential survival based on changes in stage at diagnosis and changes in treatment. For prostate cancer, there is potentially a long period between screen-detectable prostate cancer and clinical diagnosis. The lead-time between a screen-detected diagnosis and clinical diagnosis is of the order of 10 years [25, 26]. We represent the effect of screening on survival *S* as a function of time *t* = *a* – *a*_*c*_ from the possibly counter-factual age of clinical diagnosis rather than the age of screen-detected diagnosis *a*_*s*_, where we can assume that

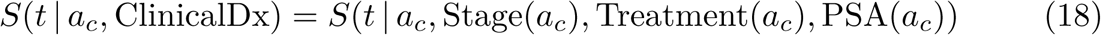

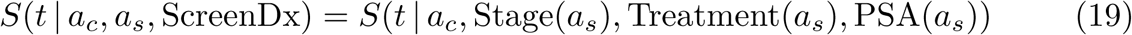

where ClinicalDx and ScreenDx represents either a clinical diagnosis or a screen-detected diagnosis, respectively, and Treatment(*a*), Stage(*a*) PSA(*a*) are the treatment modality, stage and PSA value at age *a*, respectively. The treatment sub-model assumes that the hazard ratio from SPCG-4 [27, 0.56] applies comparing both radical prostatectomy and radiation therapy with either watchful waiting or active surveillance. This point estimate is consistent with the point estimate from the PIVOT trial [28], albeit without the latter being significantly different from one. We then model survival as

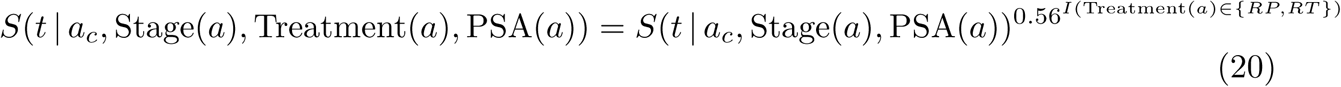

### Implementation of the Stockholm Prostata model

The FHCRC prostate cancer model was implemented in C under an open source GPL licence, although the code has not been widely distributed. We have implemented our extended model using existing C++ simulation libraries and to manage input parameters and output predictions using R. The model is implemented together with an extensible microsimulation framework. It is availablefromhttps://github.com/mclements/microsimulationandhttps://github.com/mclements/prostata under a GPL3 license, allowing for use and reuse, in contrast to most existing microsimulation models which are not open source [29].

### Model fitting and calibration

To adapt the extended prostate cancer model to the Swedish context, a number of input parameters were estimated from external data and a smaller set of parameters were estimated using simulation predictions fitted to calibration targets.

First, a set of parameters were estimated from available data sources, including parameters describing the longitudinal development of PSA with age, the rate controlling time to cancer onset, and transition between cancer states, as indicated in the first 2 rows of Table 1.

Observed characteristics of the modelled population were collected and used to estimate another set of parameters via simulations of the model. The characteristics used as calibration targets were the distribution of Gleason scores in the population, survival across age groups, and disease stage, as well as the PSA uptake (second column of Table 1). The PSA test uptake prior to 2003 was reconstructed by fitting a model to prostate cancer incidence using later PSA test rates as covariates, and we used survival analysis to estimate re-testing rates prior to cancer diagnosis by age and PSA value categories.

The model was validated against the population of Sweden and the population of Stockholm [30] (the bottom rows in Table 1). For the validation we simulated the observed PSA testing pattern and validated the model against the population data for incidence, all-cause mortality and prostate cancer mortality (results provided in the S1 Appendix).

#### Emulating the ERSPC trial

We performed a simulation experiment to emulate the ERSPC trial, where we predicted both the “control” arm and the “screening” arm with 100 million simulated men. Both arms where constructed as flat populations with inclusion between ages 55–69 years after which they where followed for 13 years. For study eligibility, we assumed that the men had not had a prostate cancer diagnosis prior to age 55 years. For the control arm, we assumed no screening. For the screening arm, we assumed four-yearly screening between ages 55 and 69 years. The PSA threshold was assumed to be 3.0 ng/mL, although in fact this varied by study site. We also used the reported biopsy compliance of 85.6%. Treatment and other-cause deaths were assumed to be similar to those observed in Stockholm.

### Calibration methods

We used four sets of targets for our calibration procedure:

1. The relative distribution of cancer staging for contemporary Sweden
2. An equality constraint on the mean time from onset to metastatic cancer
3. The incidence rate ratio due to screening from the ERSPC
4. Detailed prostate cancer survival for contemporary Sweden.

For the first step, we calibrated for the incidence-related targets 1, 2 and 3 in one likelihood; and then, as a second step, we calibrated for target 4. For targets 1, 2 and 4, we simulated for current PSA testing in Sweden; for target 3, we simulated for both arms of the ERSPC.

For target 1, we used a multinomial likelihood with unknown parameters ***θ*** = (*β*_7_, *β*_8_, *γ*_*t*_, *γ*_*m*_, *ϕ*_lag_)′. The multinomial log-likelihood was defined as

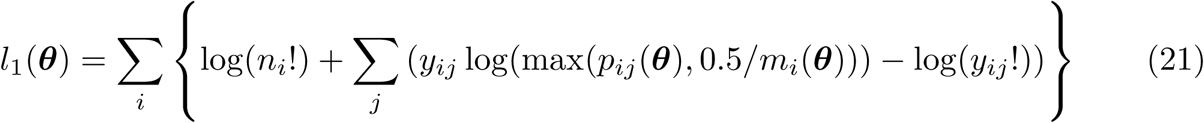

where *i* is an index over age, *j* is an index over cancer staging, *n*_*i*_ is the observed total count in a particular age group, *y*_*ij*_ is the observed count for a combination of age and cancer staging, *m*_*i*_(***θ***) is the simulated total count, and *p*_*ij*_(***θ***) is the simulated proportion of individuals in a particular disease state (see Equations (2) – (4) for the multinomial data generating mechanism). The cancer staging for the observed frequencies and simulated proportions were by age and (i) loco-regional cancers by combinations of Gleason score and T-stage, and (ii) metastatic prostate cancers. The intercept terms *α*_7_ and *α*_8_ for the distribution of Gleason score at age 35 years were not identifiable and we assumed that *α*_7_ = log(0.2) and *α*_8_=log(0.002). Half-cell corrections were performed to handle empty cells in the simulated proportions.

For target 2, we used a non-linear equality constraint on the expected time from onset to metastatic cancer to ensure identifiability of progression across T-stages. Formally,

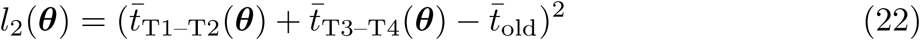

where 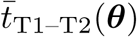 are the mean simulated transition times from onset to T3–T4, 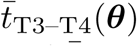 are the mean simulated transition times from T3–T4 to metastatic cancer, and 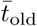 is the expected mean time from onset to metastatic cancer from a model without separate T stages (25.9 years from the FHCRC model; [12]).

For target 3, we used a non-linear equality constraint on the simulated incidence rate ratio from the ERSPC, where

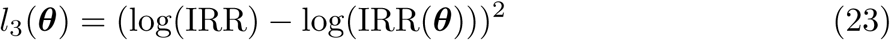

where IRR is the observed PSA screening incidence rate ratio from the ERSPC study and IRR(***θ***) is the simulated incidence rate ratio for the emulation of the ERSPC study.

Formally, the log-likelihood *l*_123_(***θ***) for targets 1–3 was

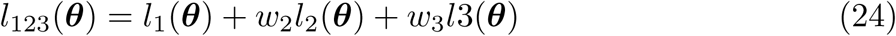

where *w*_2_ and *w*_3_ are weights for the non-linear constraints. Note that the equality constraints in targets 2 and 3 were formulated in terms of weighted quadratic penalties. The weights were selected so that the constraints were approximately satisfied (*w*_2_ = 1; *w*_3_ = 10^4^).

To optimise the simulation log-likelihood *l*_123_(***θ***), we used the Nelder-Mead optimisation algorithm. For each iteration of the optimisation, we evaluated the log-likelihood by simulating three different scenarios that depended on the parameters ***θ***. From these scenarios, we predicted values that were used in the log-likelihood, including the relative distribution of cancer staging, the mean time from onset to metastatic cancer, and the PSA screening incidence rate ratio for the reconstructed ERSPC trial. The Nelder-Mead algorithm is commonly used to optimise functions for which derivatives are difficult to calculate and for objectives that are not smooth. The standard errors were calculated from the inverse of the Hessian matrix for the negative log-likelihood (see Table 2). Given the simulation likelihood, the calculation of the Hessian matrix required that the step size for the finite differences used a larger step size (0.01).

For the second step and target 4, the distributions of Gleason score, T-stage and metastatic cancer (*θ*) were kept fixed. Using the mean between the observed 10- and 15-year survival as the calibration target and Kaplan-Meier estimates based on the model simulations, we calculated the hazard ratios by age group, cancer stage, Gleason score and PSA values. The adjustment 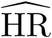.was calculated by averaging on the log hazard ratio scale

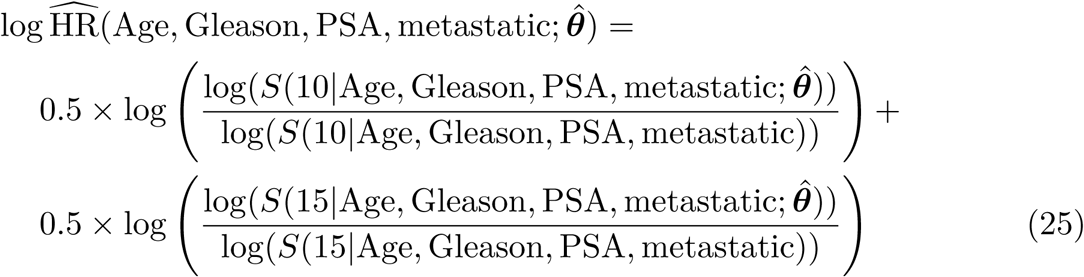

where *S*(*t |* Age, Gleason, PSA, metastatic; 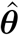) is the simulated survival to time *t* based on the parameters from step 1 and *S*(*t |* Age, Gleason, PSA, metastatic) is observed survival to time *t* from the NPCR.

#### Validation of the ERSPC mortality rate ratio

To validate against the ERSPC screening mortality RR of 0.79 (95% CI 0.61–0.88), we performed a simulation experiment to emulate the ERSPC trial (see Emulating the ERSPC trial). The mortality hazard ratio comparing the screening arm with the control arm was estimated using Poisson regression taking account of the number of prostate cancer deaths and the person-time by one-year age groups. Our validation predictions resulted in a mortality RR of 0.784 (95% Monte Carlo interval (MCI) 0.781–0.786).

## Supporting information

**S1 Appendix.** The appendix contains further detail on the model and the model inputs. It also holds a comparison of survival from diagnosis by Gleason with the FHCRC model. Finally, it also includes further validation of the model.

